# Anaerobic gut fungal communities in marsupial hosts

**DOI:** 10.1101/2023.05.31.543067

**Authors:** Adrienne L. Jones, Carrie J. Pratt, Casey H. Meili, Rochelle M. Soo, Philip Hugenholtz, Mostafa S. Elshahed, Noha H. Youssef

## Abstract

The anaerobic gut fungi (AGF) inhabit the alimentary tracts of herbivores. In contrast to placental mammals, information regarding the identity, diversity, and community structure of AGF in marsupials is extremely sparse. Here, we characterized AGF communities in sixty one fecal samples from ten marsupial species belonging to four families in the order *Diprotodontia*: *Vombatidae* (wombats), *Phascolarctidae* (koalas), *Phalangeridae* (possums), and *Macropodidae* (kangaroos, wallabies, and pademelons). Amplicon-based diversity survey using the D2 region in the large ribosomal subunit (D2 LSU) as a phylogenetic marker indicated that marsupial AGF communities were dominated by eight genera commonly encountered in placental herbivores (*Neocallimastix*, *Caecomyces*, *Cyllamyces*, *Anaeromyces*, *Orpinomyces*, *Piromyces*, *Pecoramyces*, and *Khoyollomyces*). Community structure analysis revealed a high level of stochasticity, and ordination approaches did not reveal a significant role for animal host, gut type, dietary preferences, or lifestyle in structuring marsupial AGF communities. Marsupial foregut and hindgut communities displayed diversity and community structure patterns comparable to AGF communities typically encountered in placental foregut hosts, while exhibiting a higher level of diversity and a distinct community structure compared to placental hindgut communities. Quantification of AGF load using quantitative PCR indicated a significantly smaller load in marsupial hosts compared to their placental counterparts. Isolation efforts were only successful from a single red kangaroo fecal sample and yielded a *Khoyollomyces ramosus* isolate closely related to strains previously isolated from placental hosts. Our results suggest that AGF communities in marsupials are in low abundance, and show little signs of selection based on ecological and evolutionary factors. The observed lack of host-fungal coevolutionary signal suggests a potential recent acquisition and/or a transient nature of AGF communities in marsupial herbivores.

## Introduction

Marsupials (infraclass *Marsupialia*) are mammals characterized by giving birth to undeveloped offspring and caring for them in pouches. Marsupials represent the only extant group of metatherian mammals and are endemic to Australia, North, and South America. Extant marsupials include herbivores (order *Diprotodontia*), carnivores (order *Dasyuromorphia*), and omnivores (orders *Didelphimorphia* and *Peramelemorphia*). The majority of marsupial herbivores, with rare exceptions such as the woolly opossum (genus *Caluromys*), are native to Australia and belong to the order *Diprotodontia*. Marsupial herbivores display a wide range of dietary preferences including browsers (feeding on trees and shrubs of high-growing plants, some of which can display high preference for leaves, i.e. folivores, or fruits, i.e. frugivores), grazers (feeding on grass and low-growing vegetation), and mixed feeders (Melzer et al., 2014; Arman and Prideaux, 2015; Chong et al., 2020) (Table 1).

**Table 1.** Marsupials sampled in this study, with a description of their families, species, gut type, nutritional type and habitat.

| Animal | Family | Species | Gut Type | Nutritional type | Habitat | Number of animals/habitat |
| --- | --- | --- | --- | --- | --- | --- |
| Eastern grey kangaroo | Macropodidae | Macropus giganteus | Foregut | Grazer | Sanctuary, Australia | 4 |
| Red kangaroo |  | Osphranter rufus | Foregut | Grazer | Zoo, USA | 2 |
|  |  |  |  |  | Sanctuary, Australia | 7 |
| Red-legged pademelon |  | Thylogale stigmatica | Foregut | Mixed feeder | Sanctuary, Australia | 1 |
| Red-necked wallaby |  | Notamacropus rufogriseus | Foregut | Mixed feeder | Zoo, USA | 3 |
|  |  |  |  |  | Sanctuary, Australia | 3 |
| Common brushtail possum | Phalangeridae | Trichosurus vulpecula | Hindgut | Mixed feeder | Sanctuary, Australia | 1 |
| Koala | Phascolarctidae | Phascolarctos cinereus | Hindgut | Folivore | Sanctuary, Australia | 30 |
|  |  |  |  |  | Zoo, Australia | 1 |
| Common wombat | Vombatidae | Vombatus ursinus | Hindgut | Grazer | Sanctuary, Australia | 3 |
|  |  |  |  |  | Zoo, Australia | 1 |
| Southern hairy-nosed wombat |  | Lasiorhinus latifrons | Hindgut | Grazer | Sanctuary, Australia | 4 |
|  |  |  |  |  | Zoo, Australia | 1 |

Similar to placental mammals (infraclass *Placentalia*), marsupial herbivores rely on microorganisms in their gastrointestinal tract for plant digestion and conversion to absorbable fermentation end products (Hume, 1989; Chong et al., 2020). In both groups, fermentation occurs in specialized chambers with extended food retention times to enable colonization, plant polymer mobilization and breakdown, and monomer/oligomer fermentation to soluble end products by the resident microbiota. However, herbivorous marsupial and placental guts are structurally distinct. Marsupial foregut fermenters (members of the family *Macropodidae*, e.g. kangaroos, wallabies, wallaroos, and pademelons) possess an enlarged forestomach region divided into an anterior sacciform and posterior tubiform, with fermentation processes occurring in both regions (Hume, 1989). In contrast, the majority of fermentation processes in placental foregut fermenters occurs in the rumen, a pregastric chamber that represents part of a complex four-chambered stomach. Hindgut marsupial fermenters have an enlargement of a variety of intestinal region(s), with some possessing an enlarged colon (e.g. wombats), caecum (e.g. possums), or both colon and caecum (e.g. koalas). Further, marsupial herbivores in general have a relatively lower basal metabolic rate and display an ability to forage on poor nutritional diets compared to placental mammals; adaptations seen as necessary for survival in low productive and arid habitats and a highly variable climate (Freeman, 2018).

Multiple studies have investigated microbial communities in various marsupials using culture-based (Ouwerkerk et al., 2005; Singh et al., 2015; Dhakal et al., 2020), amplicon-based (Barker et al., 2013; Gulino et al., 2013; Eisenhofer et al., 2023), and –omics-based approaches (Pope et al., 2010; Shiffman et al., 2017; Brice et al., 2019; Boath et al., 2020; Blyton et al., 2022). These studies have identified prevalent bacterial lineages in the gut of various herbivorous marsupial taxa and yielded valuable insights into the impact of ecological and evolutionary factors in shaping marsupial gut bacterial communities. However, while we have a baseline of knowledge concerning the bacterial and archaeal components of the marsupial gut, the prevalence, identity, and community structure of anaerobic gut fungi (AGF) is currently unclear. The AGF belong to a distinct basal fungal phylum (*Neocallimastigomycota*) (Li et al., 2021) and were discovered in the rumen of sheep in 1975 (Orpin, 1975). They were subsequently shown to be key constituents of the gut microbiomes of a wide range of placental mammalian herbivores (Gruninger et al., 2014). As previously noted (Gruninger et al., 2014), establishment of AGF in the gut of a herbivorous host requires long retention times and a dedicated digestive chamber (e.g. rumen, forestomach, or caecum), criteria that are satisfied in herbivorous marsupials. However, our current knowledge regarding AGF communities in marsupial herbivores is extremely sparse. An earlier review alluded to unpublished efforts pertaining to the isolation of AGF from a red kangaroo (*Macropus rufus*) (Gordon and Phillips, 1998). The isolates were putatively identified as *Piromyces* species based on microscopic observations, although diagnostic features of the genus (monocentric thalli, filamentous hyphae, and monoflagellated zoospores) have since been observed in 13 additional genera (Callaghan et al., 2015; Dagar et al., 2015; Hanafy et al., 2020b; Hanafy et al., 2022). Another review also reported on unpublished efforts where AGF rhizoidal growth was observed on plant fragments from the stomachs of four macropod species: grey kangaroo (*Macropus giganticus*), red-necked wallaby (*Macropus rufogriseus*), wallaroo (*Macropus robustus*) and swamp wallaby (*Wallabia bicolor*) (Bauchop, 1989). In addition, two previous culture-independent amplicon surveys examined AGF communities in zoo-housed red kangaroo and white-fronted wallaby (*Osphranter rufus* and *Macropus parma*) and reported a diverse community affiliated with the genera *Piromyces*, *Anaeromyces*, *Khoyollomyces*, as well as multiple yet-uncultured genera representing the most abundant AGF genera (Liggenstoffer et al., 2010).

Here we sought to characterize AGF in marsupial herbivores (order *Diprotodontia*) using culture-independent diversity surveys, qPCR quantification, and enrichment and isolation procedures. We hypothesized that the distinct gut architecture and dietary preferences of marsupial herbivores, as well as their unique evolutionary history and geographic range restriction, could select for an AGF community characterized by a high proportion of novel taxa or distinct community structure patterns compared to those of placental mammals. Surprisingly, our results suggest that the AGF communities in marsupials are neither novel nor unique. Rather, AGF appear to be present in relatively small loads or absent in marsupial gut, in contrast to their ubiquity and higher loads in placental herbivores. Further, AGF communities in marsupials appear to exhibit diversity and community structure patterns similar to those encountered in placental foregut fermenters. The ecological and evolutionary factors underpinning such observed patterns are discussed.

## Materials and Methods

### Samples

A total of 184 marsupial fecal samples were examined in this study. The samples included representatives of six different families (*Macropodidae*, *Phascolarctidae*, *Phalangeridae*, *Petauridae*, *Pseudocheiridae*, and *Vombatidae)*, fifteen different genera, and twenty different species in the order *Diprotodontia* (Table S1). These hosts encompass multiple different gut types (foregut fermenters, hindgut fermenters with an enlarged colon, caecum, or both), dietary classifications (browsers, grazers, mixed feeders), and lifestyles (zoo-housed and sanctuary-housed).

### DNA extraction and amplification

DNA extraction was conducted using a DNeasy Plant Pro kit (Qiagen®, Germantown, Maryland, USA) according to manufacturer’s instructions. The kit has previously been evaluated and utilized by multiple laboratories in prior AGF diversity surveys (Hanafy et al., 2020a; Meili et al., 2022; Young et al., 2022). Amplification of the D2 region of the large ribosomal subunit (D2 LSU) was achieved using primer pair AGF-LSU-EnvS primer pair (AGF-LSU-EnvS For: 5’-GCGTTTRRCACCASTGTTGTT-3’, AGF-LSU-EnvS Rev: 5’-GTCAACATCCTAAGYGTAGGTA-3’) (Meili et al., 2022; Young et al., 2022) modified to include the Illumina overhang adaptors. The large ribosomal subunit has been shown to be superior to ITS1, commonly used for diversity surveys of other fungal lineages, as it exhibits a much lower level of length and sequence divergence heterogeneity (Edwards et al., 2019; Hanafy et al., 2020a) and is currently the standard phylomarker in diversity surveys of AGF (Hanafy et al., 2020a; Meili et al., 2022; Young et al., 2022). PCR reactions contained 2 µl of DNA, 25 µl of the DreamTaq 2X master mix (Life Technologies, Carlsbad, California, USA), and 2 µl of each primer (10 µM) in a 50 µl reaction mix. The PCR protocol consisted of an initial denaturation for 5 min at 95°C followed by 40 cycles of denaturation at 95°C for 1 min, annealing at 55°C for 1 min and elongation at 72°C for 1 min, and a final extension of 72°C for 10 min. For samples showing negative PCR amplification in initial attempts, additional efforts were conducted (varying the DNA concentrations) and samples were only deemed negative after four attempts. Negative (reagents only) controls were included with all PCR amplifications to guard against possible cross-contamination.

### Sequencing and sequence processing

PCR products were individually cleaned using PureLink® gel extraction kit (Life Technologies, Carlsbad, California, USA) and indexed using Nexterra XT index kit v2 (Illumina Inc., San Diego, California, USA). Libraries were pooled using the Illumina library pooling calculator (https://support.illumina.com/help/pooling-calculator/pooling-calculator.htm) and pooled libraries were sequenced at the University of Oklahoma Clinical Genomics Facility (Oklahoma City, Oklahoma, USA) using the MiSeq platform. Forward and reverse Illumina reads were assembled using the make.contigs command in mothur (Schloss et al., 2009), followed by removing sequences with ambiguous bases, homopolymer stretches longer than 8 bases, and sequences that were shorter than 200 or longer than 380 bp. A two-tier approach, as detailed before (Hanafy et al., 2020a; Meili et al., 2022), was used to confidently assign sequences to either an existing cultured or uncultured genus or to a novel genus. These genus-level assignments were used to build a shared file using the mothur commands phylotype and make.shared, and the shared file was subsequently utilized as an input for downstream analysis.

### Alpha diversity measures

Alpha diversity estimates (Shannon, Simpson, and Inverse Simpson diversity indices) were calculated using the command estimate_richness in the phyloseq R package. To evaluate the importance of various factors in shaping alpha diversity patterns, only samples with at least 4 replicates at any of these factor levels were included. Comparisons were conducted between *Macropodidae*, *Phascolarctidae*, and *Vombatidae* (for host family comparison); red kangaroo, eastern grey kangaroo, koala, red-necked wallaby, southern hairy nosed wombat, and common wombat (for the animal species comparison); foregut and hindgut (for the gut type factor comparison); sanctuary and zoo (for habitat comparison), and grazer, foliovore, and mixed-feeder (for nutritional preferences comparisons). ANOVA (calculated using the aov command in R) followed by post hoc Tukey HSD tests (using TukeyHSD command in R) was used for multiple comparisons of means to identify the pairs of groups that are significantly different for each host factor.

### Community assembly and stochasticity

Two approaches were utilized to examine the contribution of various deterministic and stochastic processes in shaping community assembly: the normalized stochasticity ratio (NST) (Ning et al., 2019), and the null-model-based quantitative framework (implemented by (Stegen et al., 2015; Zhou and Ning, 2017)). The normalized stochasticity ratio was calculated using the NST package in R based on two taxonomic β-diversity dissimilarity metrics; the incidence-based Jaccard index and the abundance-based Bray-Curtis index. The function nst.boot in the NST package in R was then used to randomly draw samples within each comparison group, followed by bootstrapping of NST values. The values obtained were then compared using Wilcoxon test with Benjamini Hochberg adjustment. The iCAMP R package was used to calculate values of beta net relatedness index (βNRI) and modified Raup-Crick metric based on Bray-Curtis metric (RC_Bray_) using the function bNRIn.p. Values of βNRI and RC_Bray_ were used to partition selective processes into homogenous and heterogenous selection and stochastic processes into dispersal and drift as detailed before (Meili et al., 2022). The percentages of pairwise comparisons falling into each category were used as a proxy for the contribution of each of these processes (homogenous selection, heterogenous selection, homogenizing dispersal, and drift) to the total AGF community assembly.

### Community structure

Weighted Unifrac index was calculated using the ordinate command in the phyloseq R package and the pairwise values were used to construct PCoA ordination plots using the function plot_ordination in the phyloseq R package. PERMANOVA tests were run using the command adonis in the vegan R package. The F-statistics p-values were compared to identify factors that significantly affect the AGF community structure. The percentage variance explained by each factor was calculated as the percentage of the sum of squares of each factor to the total sum of squares.

### Comparison of marsupial and placental AGF communities

A dataset of placental mammals comprising 25 cattle, 25 goats, 25 sheep, 20 horses, 7 elephants, 3 rhinoceroses, and 3 zebras was used to compare AGF diversity and community structure patterns in marsupials to their placental counterparts (Table S2). These samples represent a fraction of samples included in a recent study of the placental AGF mycobiome (Meili et al., 2022). The dataset was combined with the 61 marsupial samples reported here and the mixed dataset was analyzed for AGF alpha diversity and community structure as described above.

### Quantitative PCR

Compared to unicellular bacteria and archaea, quantification of AGF in the herbivorous gut is problematic, given their complex life cycle (resulting in the coexistence of various stages at any given sampling time), wide variations in thallus morphology between genera (high biomass filamentous growth versus smaller bulbous thalli), drastic changes in AGF biomass pre- and post-feeding (with extremely low levels pre-feeding and a significant rapid increase post-feeding), and wide variability in DNA content/ biomass unit between polycentric and monocentric AGF taxa. Prior studies have quantified AGF load using spore counting (Fonty et al., 1987), most probable number assay (thallus forming unit) (Theodorou et al., 1990), as well as protein content, chitin content, and chitin synthase activity in cell-free extract (Gay, 1991). In addition, DNA-based qPCR methods have previously been used to quantify AGF using their SSU rRNA (Dollhofer et al., 2016), their ITS1 region (Khejornsart et al., 2011; Lwin et al., 2011; Kittelmann et al., 2012; Marano et al., 2012), the more conserved 5.8S rRNA (Edwards et al., 2008), and recently the D2 region of the LSU rRNA (Young et al., 2022). DNA-based qPCR methods have the advantage of higher sensitivity and circumventing the need for culturability.

AGF loads were quantified in 43 samples (10 kangaroos, 18 koalas, 5 wallabies, 9 wombats, and one pademelon (samples color coded in red in Table S1) using qPCR. The 25 μl PCR reaction volume contained 2 μl of extracted DNA, 0.3 μM of primers AGF-LSU-EnvS primer pair (AGF-LSU-EnvS For: 5’-GCGTTTRRCACCASTGTTGTT-3’ and AGF-LSU-EnvS Rev: 5’-GTCAACATCCTAAGYGTAGGTA-3’) (Young et al., 2022), and SYBR GreenER™ qPCR SuperMix for iCycler™ (ThermoFisher, Waltham, Massachusetts, USA), and were run on a MyiQ thermocycler (Bio-Rad Laboratories, Hercules, California, USA). The amplification protocol was comprised of heating at 95°C for 8.5 min, followed by 40 cycles, with one cycle consisting of 15 sec at 95°C and 1 min at 55°C. AGF were quantified in fecal samples as the number of LSU rRNA copies/g sample. The number of copies was calculated from the standard curve obtained from running pCR 4-TOPO or pCR-XL-2-TOPO plasmid (ThermoFisher, Waltham, Massachusetts, USA) containing an insert spanning ITS1-5.8S rRNA-ITS2-D1/D2 region of 28S rRNA from a pure culture strain. In addition to marsupial samples, AGF loads were also quantified in the feces in 40 placental mammalian AGF hosts (10 cattle, 10 goats, 10 sheep, and 10 horses) for comparative purposes. ANOVA (calculated using the aov command in R) followed by post hoc Tukey HSD tests (using TukeyHSD command in R) was used for multiple comparisons of means across marsupial host families, and species, while Student t-test (calculated using emmeans_test in the R package emmeans) was used to test the significance of difference between the two marsupial gut types and between marsupial versus placental AGF loads.

### Isolation of AGF from the marsupial gut

Isolation procedures were conducted as previously described (Hanafy et al., 2020b). Isolation efforts were conducted at 35°C and 39°C using switchgrass, cellulose, or both as a substrate. In total, 40 different enrichments were set up using 18 different marsupial fecal samples. Isolation attempts were undertaken for enrichments showing positive growth using the roll tube method as described earlier (Hungate, 1969). Isolates from the one successful enrichment were maintained at 39°C and identified using PCR and sequencing of the D1/D2 LSU using NL1 and NL4 primers as previously described (Elshahed et al., 2022).

## Results

### Amplicon-based diversity survey overview

Of the 184 marsupial samples examined, only 61 yielded AGF amplicons despite repeated attempts (Table S1). PCR amplification was successful from all or some samples belonging to 8 different marsupial species and genera: red kangaroo (*Osphranter rufus*), eastern grey kangaroo (*Macropus giganteus*), koala (*Phascolarctos cinereus* subspecies *adustus*), red-legged pademelon (*Thylogale stigmatica*), common brushtail possum (*Trichosurus vulpecula*), red-necked wallaby (*Notamacropus rufogriseus)*, southern hairy-nosed wombat (*Lasiorhinus latifrons*), and common wombat (*Vombatus ursinus*) (Table S1). AGF amplification failed from all samples belonging to eleven marsupial species: northern-tail wallaby (*Onychogalea unguifera,* n=4), agile wallaby (*Notamacropus agilis,* n=3), Bennet’s wallaby (*Macropus rufogriseus,* n=2), swamp wallaby (*Wallabia bicolor,* n=2), tammar wallaby (*Notamacropus eugenii,* n=5), parma wallaby (*Notamacropus parma,* n=1), common ringtail possum (*Pseudocheirus peregrinus,* n=3), short-eared possum (*Trichosurus caninus,* n=3), Lumholtz’s tree kangaroo (*Dendrolagus lumholtzi,* n=4), squirrel glider (*Petaurus norfolcensis,* n=1), and greater glider (*Petauroides armillatus,* n=1) (Table S1).

### Community overview

A total of 174,959 Illumina sequences of the D2 LSU region were obtained (average 2,868 ± 4,397 per sample). Phylogenetic analysis of the entire dataset demonstrated that, collectively, marsupials harbor a phylogenetically diverse AGF community. Representatives of 85 of the 87 currently reported AGF genera and candidate genera (Meili et al., 2022) were encountered (Figure 1, 2, Table S3). Only one additional novel genus (NY57) was identified as a ubiquitous (40 out of 61 samples), albeit minor (relative abundance 0.03-1.27%, Table S3), component of the AGF community in marsupials. Within individual samples, a diverse, multi-genus AGF community was observed, with an average number of genera ranging between 12-78 (1-20 if only considering genera present in >1% relative abundance) (Figure 2b).

**Figure 1.**
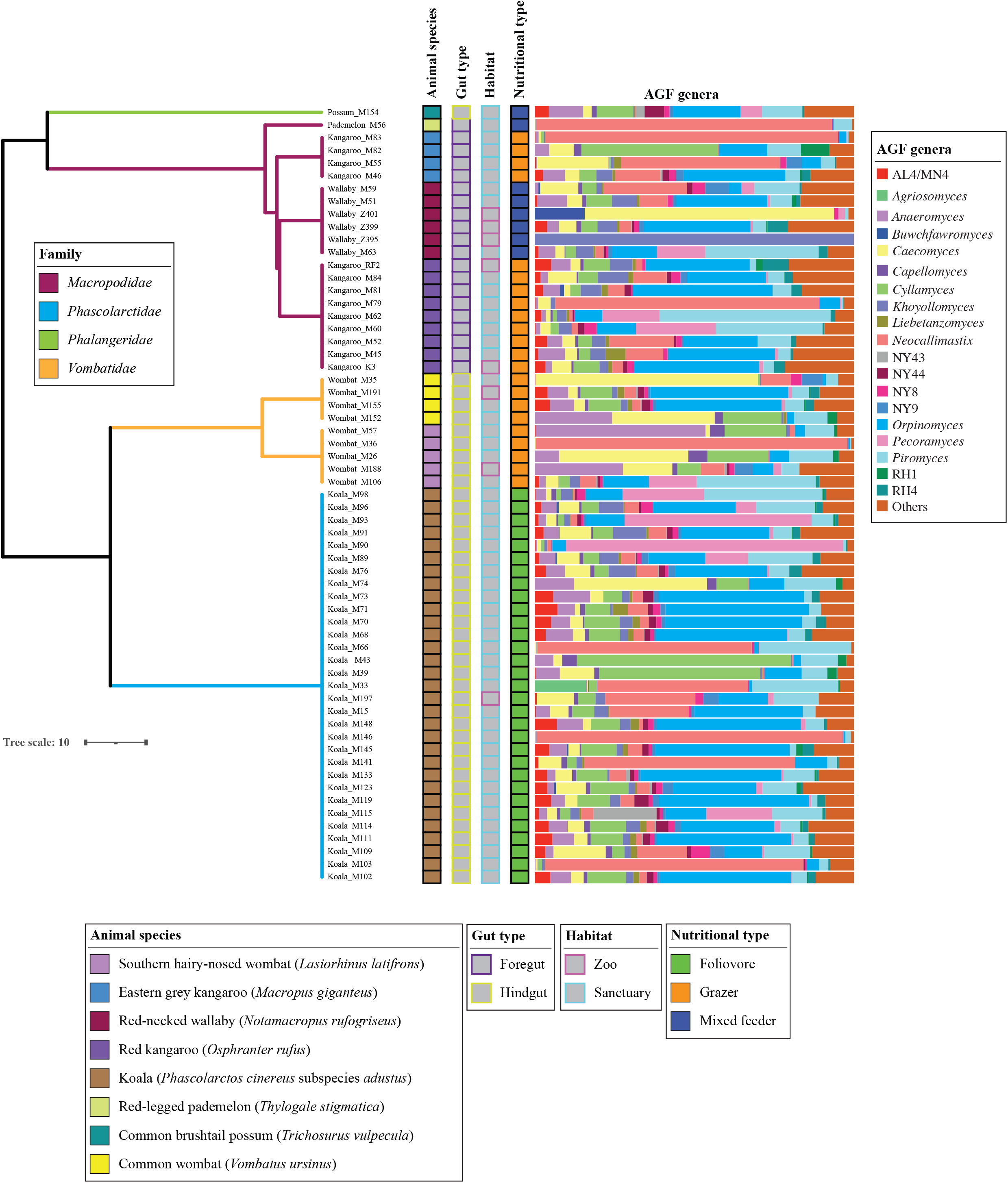
AGF community composition in the samples studied. The phylogenetic tree showing the relationship between animals was downloaded from timetree.org, and modified to include very short branch length between samples from the same animal species. Branches are color coded by family as shown in the figure legend. Tracks to the right of the tree depict the species, gut type, habitat, and nutritional type of the animals studied as shown in the figure legend. AGF genera percentage abundances are shown to the right of the tracks, with genera with <1% relative abundance grouped in “others”.

**Figure 2.**
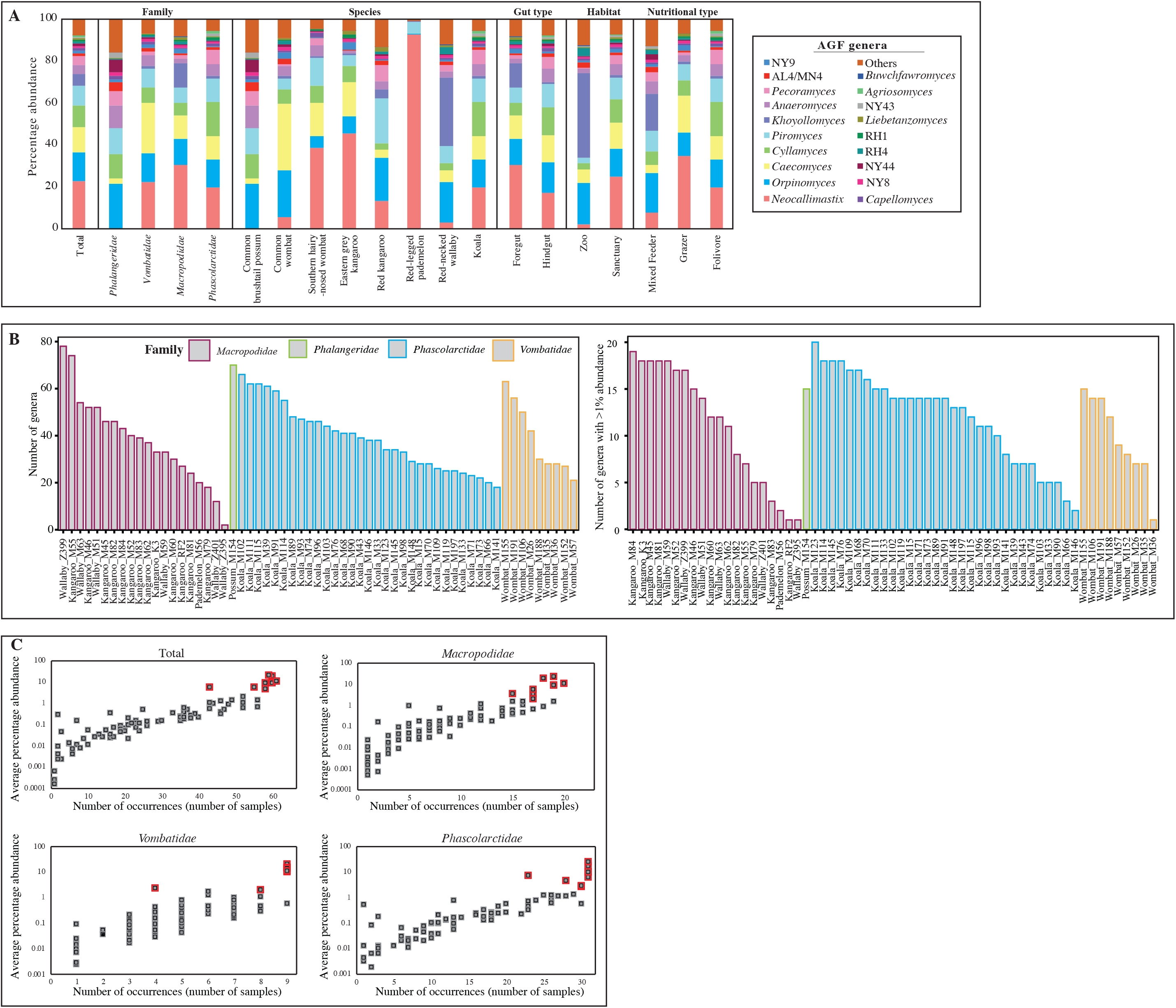
(A) AGF percentage abundance shown for all samples studied, as well as for animal families, species, gut type, habitat, and nutritional type of the animals studied. AGF genera with <1% relative abundance are grouped in “others”. (B) Total number of AGF genera (left), and number of AGF genera with >1% relative abundance (right) identified per sample. Samples names are shown on the X-axis (names match those in Figure 1), and samples are grouped by the animal family as depicted in the figure legend. (C) Relationship between occurrence (number of samples) and average relative abundance of each of the 85 genera encountered in this study. The number of samples in which the genera were identified is shown on the X-axis. Average percentage abundance across samples is plotted on the Y axis in a logarithmic scale to show genera present below 1% abundance. The 8 mostly abundant genera (*Neocallimastix*, *Orpinomyces, Caecomyces*, *Cyllamyces*, *Piromyces*, *Khoyollomyces*, *Anaeromyces*, and *Pecoramyces*) are shown with a red border. Abundance-occurrence plots are shown for all samples studied, as well as for each of the three families with > 5 animals, as depicted above each figure.

Phylogenetically, eight AGF genera represented the majority (79.33%) of the marsupial AGF communities in the entire dataset: *Orpinomyces* (19.66 ± 16.1%), *Neocallimastix* (17.23 ± 28.4%), *Piromyces* (10.04 ± 10.9%), *Caecomyces* (8.75 ± 14.77%), *Cyllamyces* (8.18 ± 11.66%), *Anaeromyces* (5.47 ± 8.05%), *Pecoramyces* (5.24 ± 14.2%) and *Khoyollomyces* (4.23 ± 12.72%) (Figure 2a). The predominance of these genera was observed across the marsupial families *Macropodidae* (75.55%), *Phascolarctidae* (85.6%), and *Vombatidae* (84.65%). In addition to their high relative abundance, these eight genera were also ubiquitous across the three families (Figure 2c, red boxes) with a positive correlation observed between relative abundance and prevalence (Figure 2c).

### Alpha diversity estimates

AGF alpha diversity patterns were assessed using three different indices: Shannon (Figure 3), Simpson, and Inverse Simpson (Figure S1). ANOVA results show a comparable level of alpha diversity between all families and species examined; as well as between foregut and hindgut fermenters, zoo- and sanctuary-housed animals, and nutritional types (Figure 3a, S1a).

**Figure 3.**
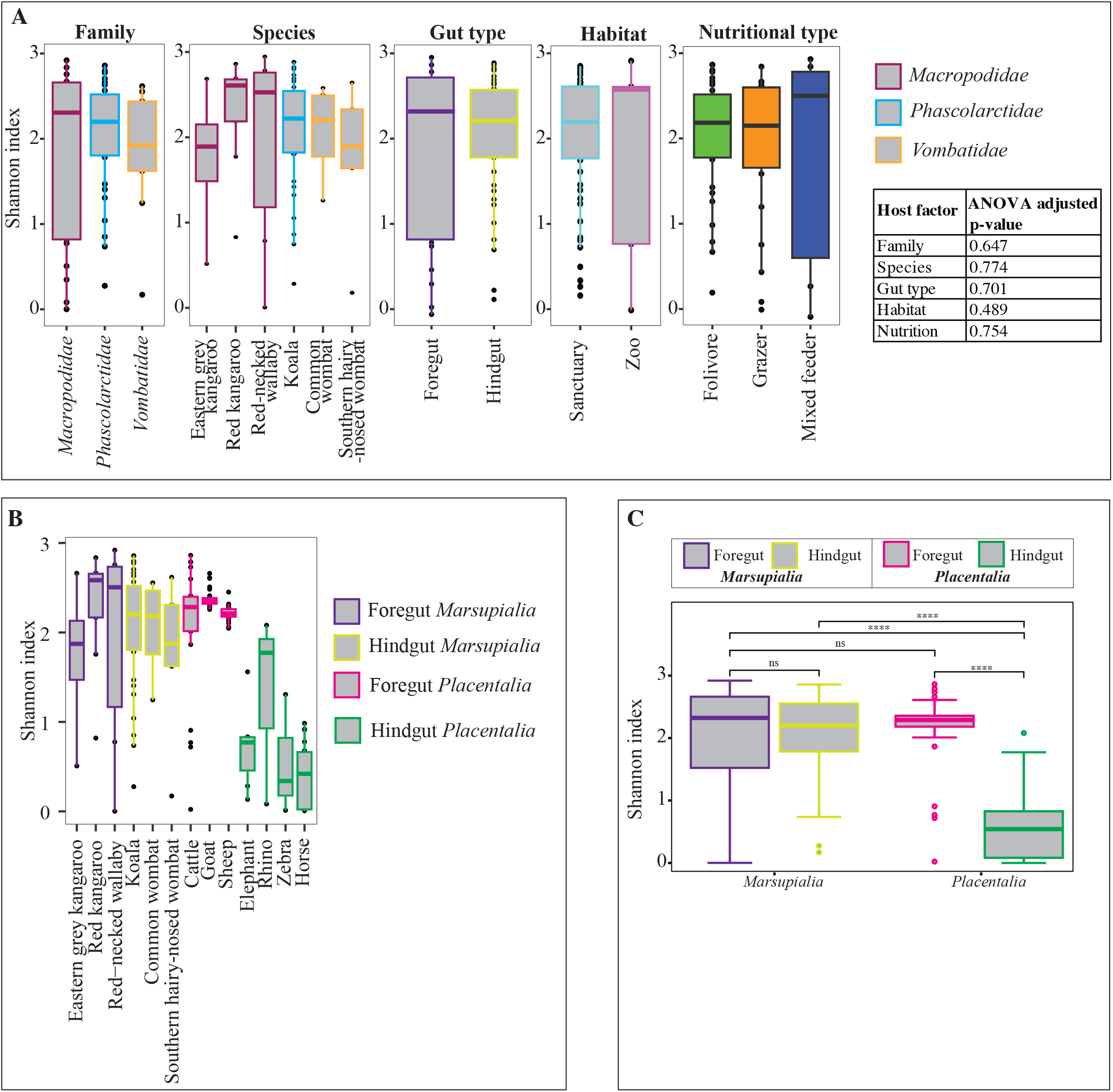
Alpha diversity patterns. (A) Box and whisker plots showing the distribution of Shannon diversity index for different families, species, gut types, habitats, and nutritional types of the animals studied. Results of ANOVA are shown in the table to the right. (B) Box and whisker plots showing the distribution of Shannon diversity index for animals species, color coded by their gut type, in comparison with foregut and hindgut placental animals representatives. (C) Results of Tukey post-hoc tests for pairwise infraclass-gut type comparisons. ns: not significant, ****: p-value <0.0001.

In addition to comparing alpha diversity patterns amongst various marsupial hosts, we also compared marsupial AGF alpha diversity patterns to their placental counterparts. The results indicated that all marsupial species (regardless of their gut type) harbor an AGF community with a comparable alpha diversity (ANOVA p-value >0.05) to placental foregut fermenters (Figure 3b, Figure S1b-c, Table S4), and a significantly higher alpha diversity than placental hindgut fermenters (Figure 3b, Figure S1b-c, Table S4).

### Stochastic processes play an important role in shaping AGF community in marsupials

Normalized stochasticity ratios (NST) indicate that, regardless of the β-diversity index used (abundance-based Bray-Curtis index, and incidence-based Jaccard index), stochastic, rather than deterministic, processes are the major contributors to AGF community assembly in marsupials (NST values of 70-75.7% for families, 51.4-90% for species, 72.5-76% for gut type, 69.9-76.4% for habitat, and 69.4-78.5% for nutritional type) (Figure 4a). AGF community assembly in the different marsupial species significantly differed in their stochasticity, with values increasing in the order: red kangaroo < southern hairy-nosed wombat < koala < common wombat < red necked wallaby < eastern grey kangaroo (Wilcoxon test p-value <0.05) (Figure 4a).

**Figure 4.**
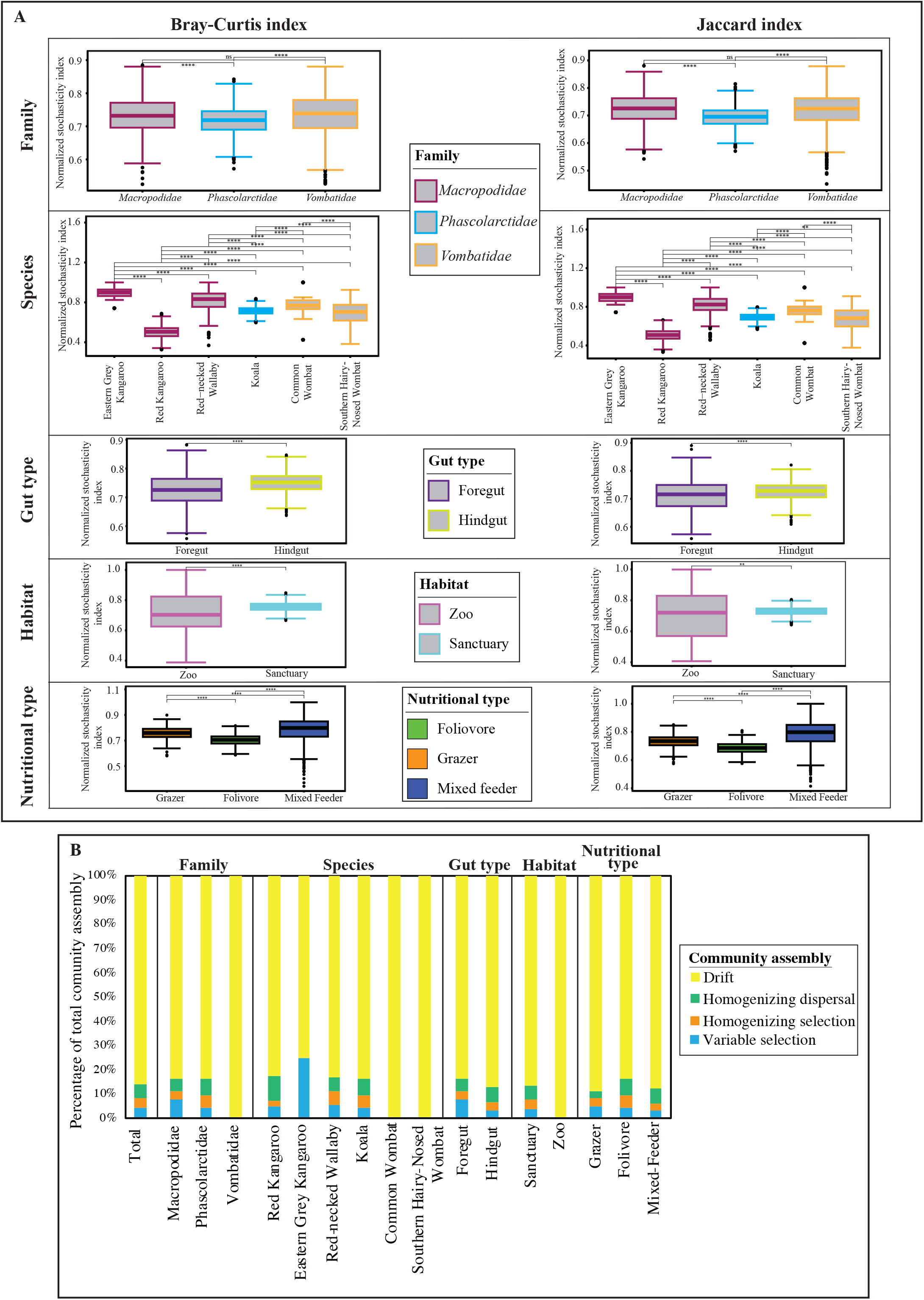
AGF community assembly in marsupial hosts. (A) Box and whisker plots showing the distribution of the bootstrapping results (n=1000) for the levels of stochasticity in AGF community assembly calculated as normalized stochasticity ratio (NST). Results compare different animal families (top row), animal species (second row), gut type (third row), habitat (fourth row), and nutritional type (fifth row). Two normalized stochasticity ratio (NST) were calculated; the abundance-based Bray-Curtis index (left), and the incidence-based Jaccard index (right). ****: Wilcoxon p-value <0.0001; ns: not significant. (B) The percentages of the various deterministic and stochastic processes shaping AGF community assembly of the total dataset, and when sub-setting for different animal families, species, gut types, habitats, and nutritional types.

To quantify the contribution of specific stochastic (homogenizing dispersal, dispersal limitation, and drift) processes in shaping the AGF community assembly in marsupials, we employed the previously suggested two-step null-model-based quantitative framework (Stegen et al., 2015; Zhou and Ning, 2017). Results (Figure 4b) broadly confirmed the patterns observed with NST values, where the AGF community assembly is mostly stochastic. The majority of stochasticity is caused by drift across all host species, families, gut types, habitats, and nutritional types examined (Figure 4b).

### Community structure patterns

AGF community structure in marsupials was assessed using PCoA based on the phylogenetic similarity-based beta diversity index weighted Unifrac. The first two axes explained 53.3% of the variance. Similar to alpha diversity estimates, the results demonstrated no clear role for marsupial family (Figure 5a, PERMANOVA p-value=0.063), species (Figure 5a, PERMANOVA p-value=0.08), gut type (Figure 5a, PERMANOVA p-value=0.298), habitat (Figure 5a, PERMANOVA p-value=0.086), or nutritional type (Figure 5a, PERMANOVA p-value=0.159) in shaping AGF community structure in marsupials.

**Figure 5.**
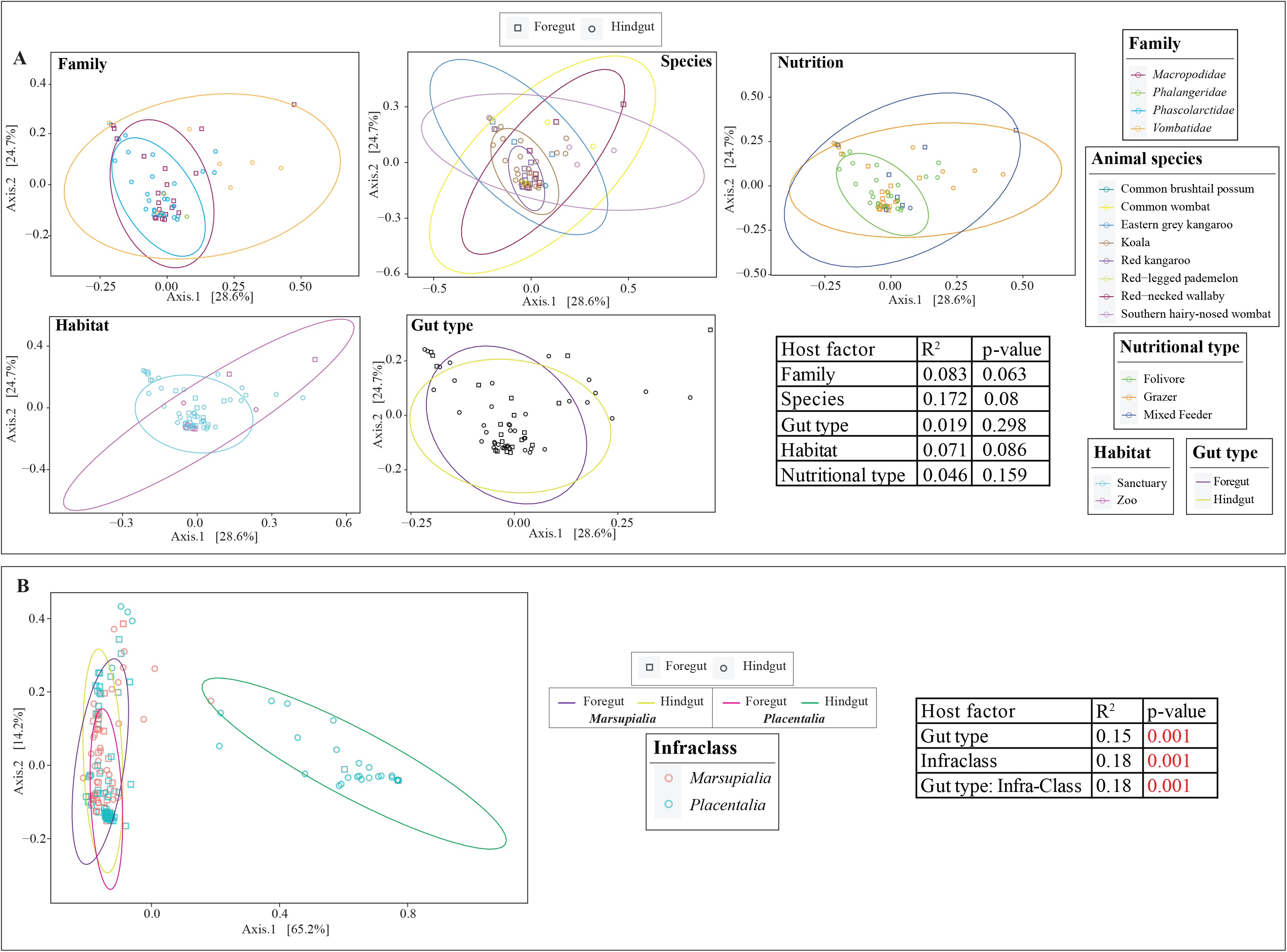
Ordination plots based on AGF community structure in the studied hosts. (A) Principal coordinate analysis (PCoA) ordination plots based on AGF community structure were constructed using the phylogenetic similarity-based weighted Unifrac index. The % variance explained by the first two axes is displayed on the axes. Samples are color coded by animal family, species, nutritional type, habitat, and gut type as shown in the legend on the right hand side, while the shape depicts the gut type as shown in the figure legend on top. Ellipses encompassing 95% of variance are shown for each of the factors and are color coded similar to the samples. Results of PERMANOVA test for partitioning the dissimilarity among the sources of variation (including animal family, species, gut type, habitat, and animal nutritional type) are shown in the table to the right. The F-statistic R^2^ depicts the fraction of variance explained by each factor, while the p-value depicts the significance of the host factor in affecting the community structure. (B) AGF community structure in marsupial hosts in comparison to a dataset of 75 placental foregut fermenters (25 cattle, 25 goats, and 25 sheep), and 33 placental hindgut fermenters (20 horses, 7 elephants, 3 rhinoceroses, and 3 zebras). Samples are color coded by their infraclass (*Marsupialia* versus *Placentalia*) and ellipses encompassing 95% of variance are shown for the four sub-categories (foregut *Marsupialia*, foregut *Placentalia*, hindgut *Marsupialia*, hindgut *Placentalia*) with color codes depicted in the figure legend to the right. Results of PERMANOVA test for partitioning the dissimilarity among host infraclass, gut type, and the interaction between the two are shown in the table to the right, where the F-statistic R^2^ depicts the fraction of variance explained by each factor, while the p-value depicts the significance of the host factor in affecting the community structure.

However, when marsupials’ AGF community structure was compared to that of placental mammals, all marsupial samples showed a clear clustering pattern close to foregut placental hosts, with hindgut placental host clustering separately (Figure 5b). To partition the dissimilarity among the sources of variation (host infraclass and gut type), we ran PERMANOVA tests (Anderson and Walsh, 2013) with the addition of interaction terms (to test for gut type specific differences in the host infraclass). Host infraclass, gut type, and the interaction of both, all significantly influenced community structure (p-value=0.001), with the largest effect being the infraclass (18% of variance), and its interaction with gut type (18% of variance).

### AGF loads in marsupial hosts

AGF load was tested in 43 samples representing the three well sampled marsupial families *Macropodidae*, *Phascolarctidae*, and *Vombatidae*, as well as 7 of the 8 marsupial species studied here (red kangaroo (n=8), eastern grey kangaroo (n=2), red-legged pademelon (n=1), red-necked wallaby (n=5), koala (n=18), common wombat (n=4), and southern hairy-nosed wombat (n=5)). AGF load in all examined marsupials was low (1.19 × 10^2^ ± 3.6 × 10^2^ copies/g feces). No significant differences were observed based on animal family (Tukey post-hoc test p-value >0.8), species (Tukey post-hoc test p-value >0.8), or gut type (Student t-test p-value =0.52) (Figure 6a-c).

**Figure 6.**
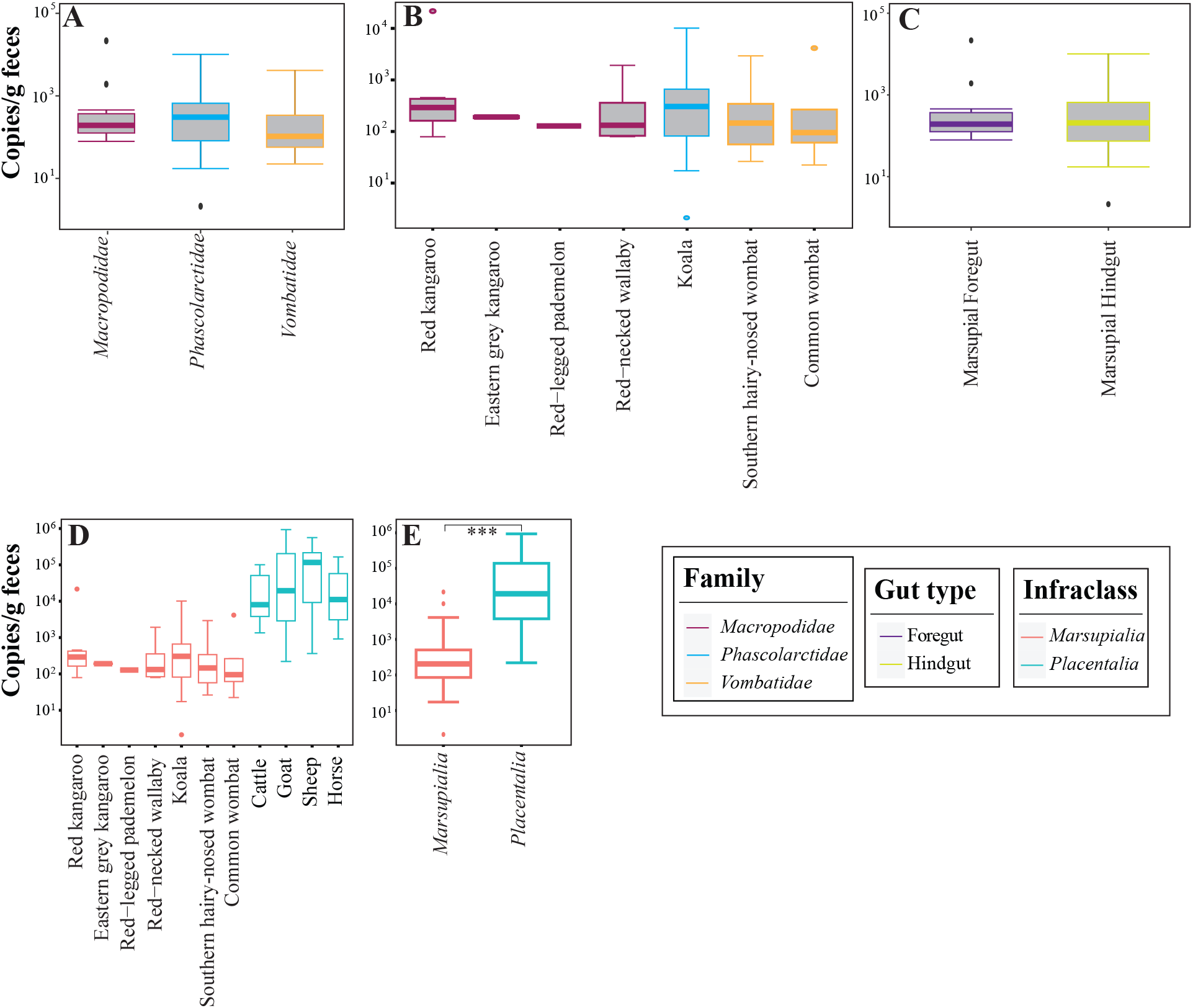
AGF load in the 43 marsupial samples examined using quantitative PCR. Boxplots showing the distribution of AGF load in the three marsupial families (A), seven marsupial species (B), and two gut types (C). (D) Comparison to AGF load in the 43 marsupial hosts to 40 placental counterparts. Boxplots in (E) show the distribution of AGF load in the 43 marsupial hosts (infra class *Marsupialia*) versus the 40 placnetal hosts (infraclass *Placentalia*). **: Student t-test p-value =0.004.

For comparison, AGF load quantified in 40 placental samples (representing 10 cattle, 10 goat, 10 sheep, and 10 horses) was significantly higher (average= 1.01 × 10^5^ ± 1.82 × 10^5^ copies/g feces) compared to marsupial mammals (Student t-test p-value =0.0005) (Figure 6d-e).

### Isolation

Attempts were made to obtain AGF isolates from freshly-collected marsupial fecal samples. Despite our best efforts, isolation was only successful from one red kangaroo sample out of forty different enrichment attempts (purple text in Table S1). Five isolates were obtained from a single red kangaroo sample (Table S1). The five isolates were identified as *Khoyollomyces ramosus* (Figure 7) and their D1/D2 LSU markers were 0.95-3.6% divergent from the *Khoyollomyces ramosus* type strain ZS33 (GenBank accession number MT085710).

**Figure 7.**
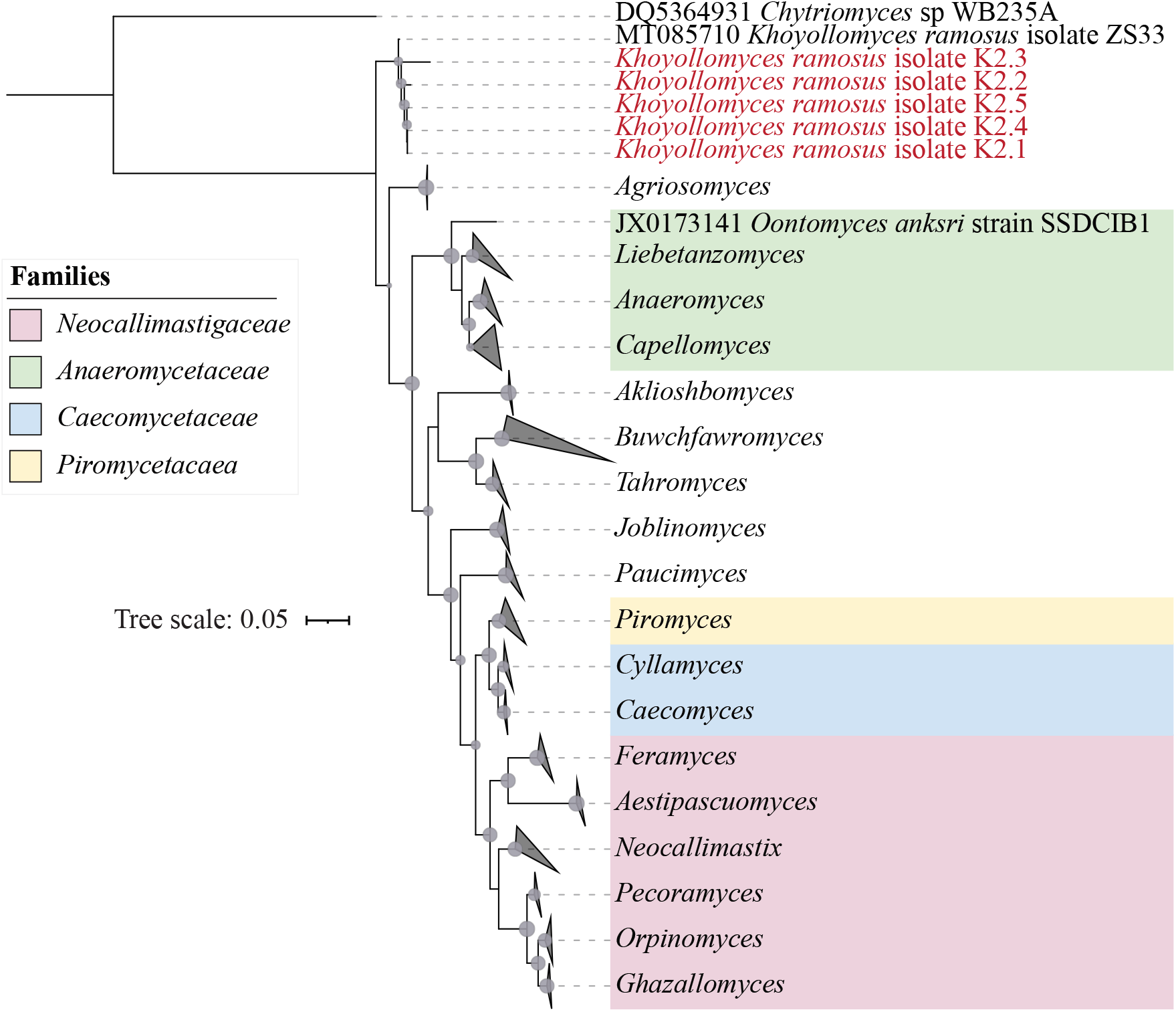
Assessment of the phylogenetic position of the newly obtained isolates from a kangaroo using D1/D2 LSU as a phylogenetic markers. The tree was constructed using the maximum likelihood approach implemented in FastTree. Scale bar indicates the number of substitutions per site. Bootstrap values are shown for nodes with >70% support as grey spheres, where the size of the sphere is proportional to the bootstrap value. The four previously suggested *Neocallimastigomycota* families are color coded as shown in the figure legend. New isolates are identified as *Khoyollomyces ramosus* and are shown in red text.

## Discussion

In this study, we investigated the AGF community in marsupial hosts. AGF occurrence was identified in 61/184 samples. The AGF communities in marsupials were dominated by genera previously identified as predominant members of the placental mammalian gut mycobiome (Meili et al., 2022) (Figures 1 and 2). Diversity and community structure patterns were comparable across all marsupial samples regardless of the animal host family, species, gut type, habitat, or nutritional classification (Figures 3 and 5). Assembly of AGF communities is predicted to be largely shaped by stochastic (mostly drift) rather than deterministic processes (Figure 4). Further, marsupial AGF communities were highly similar to those encountered in foregut, but not hindgut, placental herbivores (Figure 5). Repeated attempts to isolate AGF from marsupials yielded only five closely-related isolates that were < 3.6% divergent from the type strain of *Khoyollomyces ramosus* previously isolated from a zebra (Hanafy et al., 2020b) (Figure 7).

As described above, we hypothesized that given their unique gastrointestinal tract structures, dietary preferences, and geographic range restriction; marsupials’ AGF communities could exhibit significant differences in diversity and community structure when compared to their placental counterparts. However, our results indicate that marsupial AGF communities were neither novel nor unique, but rather dominated by well-characterized genera and candidate genera previously identified as predominant taxa in the placental gut (Meili et al., 2022). Further, given the differences in gut type (sacciform and tubiform enlarged foregut, enlarged caecum, enlarged colon, or enlarged caecum and colon), and dietary preferences (browsers, grazers, mixed feeders) between various marsupial species examined, we expected AGF communities to display clear distinctions based on ecological or host-associated factors. Again, our initial expectation was not fulfilled, with comparative levels of alpha diversity and highly similar community structure patterns encountered across all marsupial samples. Such lack of clear differences in AGF diversity and community structure between marsupials is in stark contrast to the clear host-driven stratification of AGF communities in placental mammals (Meili et al., 2022).

We argue that the failure to identify novel AGF taxa in marsupials, as well as the lack of distinct marsupial community composition patterns (e.g. communities where the majority of AGF sequences are affiliated with minor/rare members of the AGF communities in placental mammals), indicate that marsupial host evolution was not associated with a parallel process of evolution of marsupial-specific AGF lineages. However, we could not conclusively rule out the potential occurrence of such a process in extinct marsupials or in hosts not sampled in this study. Further, the lack of clear AGF community structure differences between various marsupial hosts based on ecological and evolutionary factors suggests a passive acquisition from foregut placentals. The reason behind the exclusive acquisition of AGF community in all marsupials, regardless of their gut type, from placental foregut donors remains unclear; but could possibly be attributed to the higher number of placental foregut animals (e.g. cattle, goat, sheep) compared to hindgut fermenters (e.g. horses), hence allowing higher incidences of contact and transmission through fecal exposure.

The important role played by AGF in plant biomass degradation in placental mammals has long been recognized. AGF were shown to initiate plant biomass colonization (Orpin and Bountiff, 1978; Edwards et al., 2008) and produce a wide array of highly efficient lignocellulolytic enzymes (Steenbakkers et al., 2003; Steenbakkers et al., 2008; Comlekcioglu et al., 2010; Novotná et al., 2010; O’Malley et al., 2012; Cao et al., 2013; Gruninger et al., 2014; Morrison et al., 2016a; Morrison et al., 2016b; Lange et al., 2019). However, their role and relative contribution to plant biomass degradation in marsupials remain unclear. Quantification of AGF load using qPCR showed significantly lower levels (expressed as ribosomal operon copy number/g of feces) compared to placental mammals (Figure 6). Also, PCR amplification failed in 62.5% of the samples examined and enrichment attempts were only successful in 10/32 samples. These low AGF loads, especially when coupled to the observed lack of host-selection patterns (Figure 5), high level of stochasticity (Figure 3), and apparent passive acquisition patterns from placental hosts, collectively point to a minor role for AGF in marsupial feed digestion. This could be a reflection of marsupial preference to a wider range of diets, many of which have a lower proportion of cellulose and arabinoxylan hemicellulose, the preferred substrates for AGF [19, 20]. Whether AGF abundance in marsupial gut microbiomes and their relative importance in the digestive process dynamically varies in individual subjects based on diet composition (e.g., increasing in kangaroos fed fresh grass diet, but decreasing when browsing on shrubs) remains to be seen.

Following acquisition, it is unknown how well AGF communities are retained in marsupial hosts. Also, the role and relative contribution of vertical (mother to offspring) versus horizontal (acquisition through direct contact or exposure to fecal matter of other animals) transmission in maintaining communities are largely unknown. The observed low AGF load could potentially hinder effective vertical transmission and render the community more prone to loss under adverse conditions (e.g., scarcity of diet, changes in diet composition, sickness, and dysbiosis). This could necessitate continuous horizontal transmission (through direct animal-to-animal contact or exposure to fecal matter) from other marsupial or placental subjects. Evidence of long-term survivability of AGF in dried feces, possibly through the formation of long-term survival structures (Milne et al., 1989; McGranaghan et al., 1999; Gruninger et al., 2014), has previously been reported, a trait that can facilitate cross-subject horizontal transmission in AGF. The proposed continuous need for horizontal transmission and proposed minor role for AGF in the marsupial gut, could account for our inability to detect AGF occurrence in 123 out of 184 samples examined. On the other hand, the high transmissibility of AGF could also facilitate vertical transmission aided by the close proximity associated with extended nurturing and caring of offspring in marsupials.

The geographic isolation of Australia from Gondwana occurred approximately 100 Mya, resulting in the complete separation of Australia from Antarctica (≈45 Mya) and South America (≈30 Mya) (van den Ende et al., 2017). The dominance of marsupials throughout Australia’s natural history post separation from Gondwana, and the lack of native placental mammals in Australia has been well documented (Woodburne and Case, 1996). As such, given the central role played by placental mammalian evolution in shaping AGF evolution, maintenance, and dissemination (Meili et al., 2022), the proposed lack of a parallel process in marsupials, and the proposed role of placental hosts in seeding marsupial gut microbiomes with AGF; timing the acquisition of AGF by marsupial hosts represents an interesting dilemma. The lack of historic interaction between placental and marsupial herbivores in Australia represents a bottleneck hindering AGF acquisition during the early stages of marsupial herbivores’ evolution (66 Mya) (Amador and Giannini, 2021) and subsequent evolution of the order *Diprotodontia* (53 Mya) (Meredith et al., 2008; Mitchell et al., 2014), the hindgut family *Phalangeridae* (Mid Eocene, ∼45 Mya) (Meredith et al., 2008; Mitchell et al., 2014), the split between *Vombatidae*, *Phascolarctidae* (split early Oligocene, ∼30 Mya) (Meredith et al., 2008; Mitchell et al., 2014), and the evolution of the foregut family *Macropodidae* (Mid Miocene, ∼15-18 Mya) (Meredith et al., 2008; Mitchell et al., 2014).

The only recorded instances of placental mammals arriving in Australia prior to human colonization are bats, rodents, and dugongs visiting the shores of the continents. The colonization of Australia by Aboriginal Australians (≈50,000 years ago) could represent another opportunity for placental mammalian introduction. Conversely, the colonization by European settlers, commencing in the late 1780s, has certainly led to the introduction of multiple placental mammals, including many herbivores, to Australia. Given that timeline, an earlier AGF seeding of marsupials by placentals prior to human colonization appears unlikely, given the lack of AGF in bats and rodent guts and the extreme transient nature of potential interactions between the herbivorous hindgut fermenting dugong with marsupials. As well, while we reason that Aboriginal Australians arrival to Australia has introduced some placental species (e.g. dingo), there is no concrete evidence for the widescale introduction of AGF-harboring placental herbivores during this earlier wave of human colonization. Therefore, we raise the intriguing possibility that AGF occurrence in marsupial hosts represents a very recent phenomenon, enabled by the large-scale introduction of cattle, and other large placental herbivores into Australia, post-European colonization.

In conclusion, our study is the first to provide a detailed analysis of the marsupial AGF community. We provide a thorough analysis of the patterns of occurrence, identity, loads, diversity, and community structure of AGF in marsupial hosts and use these results to provide insights on the possible role of AGF in the marsupial gut microbiome, acquisition and retention patterns of AGF in marsupials, co-evolutionary patterns, or lack thereof, between marsupials and AGF, and potential timing of AGF colonization of the marsupial gut.

## Acknowledgments

We thank Lone Pine Koala Sanctuary, Wildlife Habitat, Cairns Tropical Zoo, David Fleay Wildlife Park, D. S. Teakle and Amy Shima for collecting samples. This work was supported by the NSF grant number 2029478 to MSE and NHY, and an ARC Discovery Project (DP150104202) awarded to PH and RMS.

## Contributions

**ALJ:** Formal analysis, methodology, data curation, visualization, writing original draft.

**CJP:** Formal analysis, methodology, data curation, writing-review & Editing.

**CHM:** Formal analysis, methodology, data curation, writing-review & Editing.

**RMS:** Data curation, funding acquisition, methodology, resources, writing-review & Editing.

**PH:** Data curation, funding acquisition, methodology, resources, writing-review & Editing.

**MSE:** Conceptualization, funding acquisition, project administration, supervision, validation, visualization, writing-original draft.

**NHY:** Conceptualization, data curation, formal analysis, funding acquisition, project administration, supervision, validation, visualization, writing-review & Editing.

