## Supplementary document for "Anaerobic gut fungal communities in marsupial hosts"

**Supplementary tables:**

**Table S1.** Summary of datasets examined in this study. Samples are grouped by animal host family, then animal host species. Gut type, country of origin, habitat, and nutritional classification are also shown. The 61 samples that produced amplicons are shown on top followed by the samples where AGF amplification was unsuccessful. The 43 samples that were used for AGF quantification (using qPCR) are shown in red text, and the one sample yielding isolates is shown in purple text.

**Table S2.** Placental hosts utilized for AGF diversity (both alpha and beta diversity) and load (qPCR quantification) comparisons to the marsupial hosts studied here. The sample name column correlates to samples studied in a previous global herbivorous mycobiome analysis [27].

**Table S3.** Good's coverage and AGF genus-level community composition (shown as percentage abundance) for the datasets studied. Samples are shown in the same order as in Table S1.

**Table S4.** Tukey post-hoc adjusted p-value for the comparison of AGF alpha diversity measures (Shannon, inv-Simpson, and Simpson indices) between marsupial and placental mammals. Significant p-values (<0.05) are highlighted in red.

| Placental mammal gut type | Placental mammal species | Marsupial mammal species | Tukey Post-Hoc adjusted p-value | | |
|---|---|---|---|---|---|
| | | | Shannon | Inv-Simpson | Simpson |
| Foregut Ruminant | Cattle | Common brushtail possum | 0.992 | 0.597 | 1.000 |
| | | Common wombat | 1.000 | 0.994 | 1.000 |
| | | Eastern grey kangaroo | 0.999 | 0.962 | 0.996 |
| | | Koala | 1.000 | 0.999 | 1.000 |
| | | Red kangaroo | 1.000 | 1.000 | 1.000 |
| | | Red-legged pademelon | 0.195 | 0.628 | 0.110 |
| | | Red-necked wallaby | 1.000 | 1.000 | 0.995 |
| | | Southern hairy-nosed wombat | 0.996 | 0.990 | 0.997 |
| | Goat | Common brushtail possum | 1.000 | 0.274 | 1.000 |
| | | Common wombat | 0.999 | 1.000 | 1.000 |
| | | Eastern grey kangaroo | 0.727 | 1.000 | 0.967 |
| | | Koala | 0.765 | 1.000 | 0.999 |
| | | Red kangaroo | 1.000 | 0.966 | 1.000 |
| | | Red-legged pademelon | 0.047 | 0.905 | 0.069 |
| | | Red-necked wallaby | 0.894 | 1.000 | 0.948 |
| | | Southern hairy-nosed wombat | 0.553 | 1.000 | 0.967 |
| | Sheep | Common brushtail possum | 0.999 | 0.146 | 1.000 |
| | | Common wombat | 1.000 | 1.000 | 1.000 |
| | | Eastern grey kangaroo | 0.964 | 1.000 | 0.993 |
| | | Koala | 1.000 | 0.891 | 1.000 |
| | | Red kangaroo | 1.000 | 0.603 | 1.000 |
| | | Red-legged pademelon | 0.105 | 0.975 | 0.098 |
| | | Red-necked wallaby | 0.998 | 0.978 | 0.990 |
| | | Southern hairy-nosed wombat | 0.912 | 1.000 | 0.994 |
| Hindgut | Elephant | Common brushtail possum | 0.051 | 0.010 | 0.453 |
| | | Common wombat | 0.025 | 0.848 | 0.293 |
| | | Eastern grey kangaroo | 0.258 | 0.941 | 0.877 |
| | | Koala | 0.000 | 0.015 | 0.007 |
| | | Red kangaroo | 0.000 | 0.009 | 0.014 |

| | | | | | |
|---|---|---|---|---|---|
| | | Red-legged pademelon | 1.000 | 1.000 | 0.992 |
| | | Red-necked wallaby | 0.021 | 0.097 | 0.639 |
| | | Southern hairy-nosed wombat | 0.178 | 0.734 | 0.734 |
| | Horse | Common brushtail possum | 0.005 | 0.001 | 0.008 |
| | | Common wombat | 0.000 | 0.207 | 0.000 |
| | | Eastern grey kangaroo | 0.004 | 0.359 | 0.000 |
| | | Koala | 0.000 | 0.000 | 0.000 |
| | | Red kangaroo | 0.000 | 0.000 | 0.000 |
| | | Red-legged pademelon | 1.000 | 1.000 | 1.000 |
| | | Red-necked wallaby | 0.000 | 0.001 | 0.000 |
| | | Southern hairy-nosed wombat | 0.001 | 0.087 | 0.000 |
| | Rhino | Common brushtail possum | 0.596 | 0.108 | 0.889 |
| | | Common wombat | 0.941 | 1.000 | 0.974 |
| | | Eastern grey kangaroo | 1.000 | 1.000 | 1.000 |
| | | Koala | 0.688 | 0.926 | 0.866 |
| | | Red kangaroo | 0.451 | 0.695 | 0.758 |
| | | Red-legged pademelon | 0.981 | 1.000 | 0.925 |
| | | Red-necked wallaby | 0.978 | 0.922 | 1.000 |
| | | Southern hairy-nosed wombat | 1.000 | 1.000 | 1.000 |
| | Zebra | Common brushtail possum | 0.051 | 0.010 | 0.079 |
| | | Common wombat | 0.059 | 0.807 | 0.021 |
| | | Eastern grey kangaroo | 0.328 | 0.902 | 0.167 |
| | | Koala | 0.002 | 0.106 | 0.001 |
| | | Red kangaroo | 0.001 | 0.045 | 0.001 |
| | | Red-legged pademelon | 1.000 | 1.000 | 1.000 |
| | | Red-necked wallaby | 0.068 | 0.174 | 0.064 |
| | | Southern hairy-nosed wombat | 0.270 | 0.720 | 0.091 |

**Supplementary figures:**

**Figure S1.** (A) Box and whisker plots showing the distribution of Simpson and Inverse Simpson diversity indices for different families, species, gut types, habitats, and nutritional types of the animals studied. Results of ANOVA are shown in the table to the right. (B) Box and whisker plots showing the distribution of Simpson and Inverse Simpson diversity indices for animal species, color coded by their gut type, in comparison to foregut and hindgut placental animals representatives. (C) Results of Tukey post-hoc tests for pairwise infraclass-gut type comparisons. ns: not significant, ****: $p\text{-value}<0.0001$.

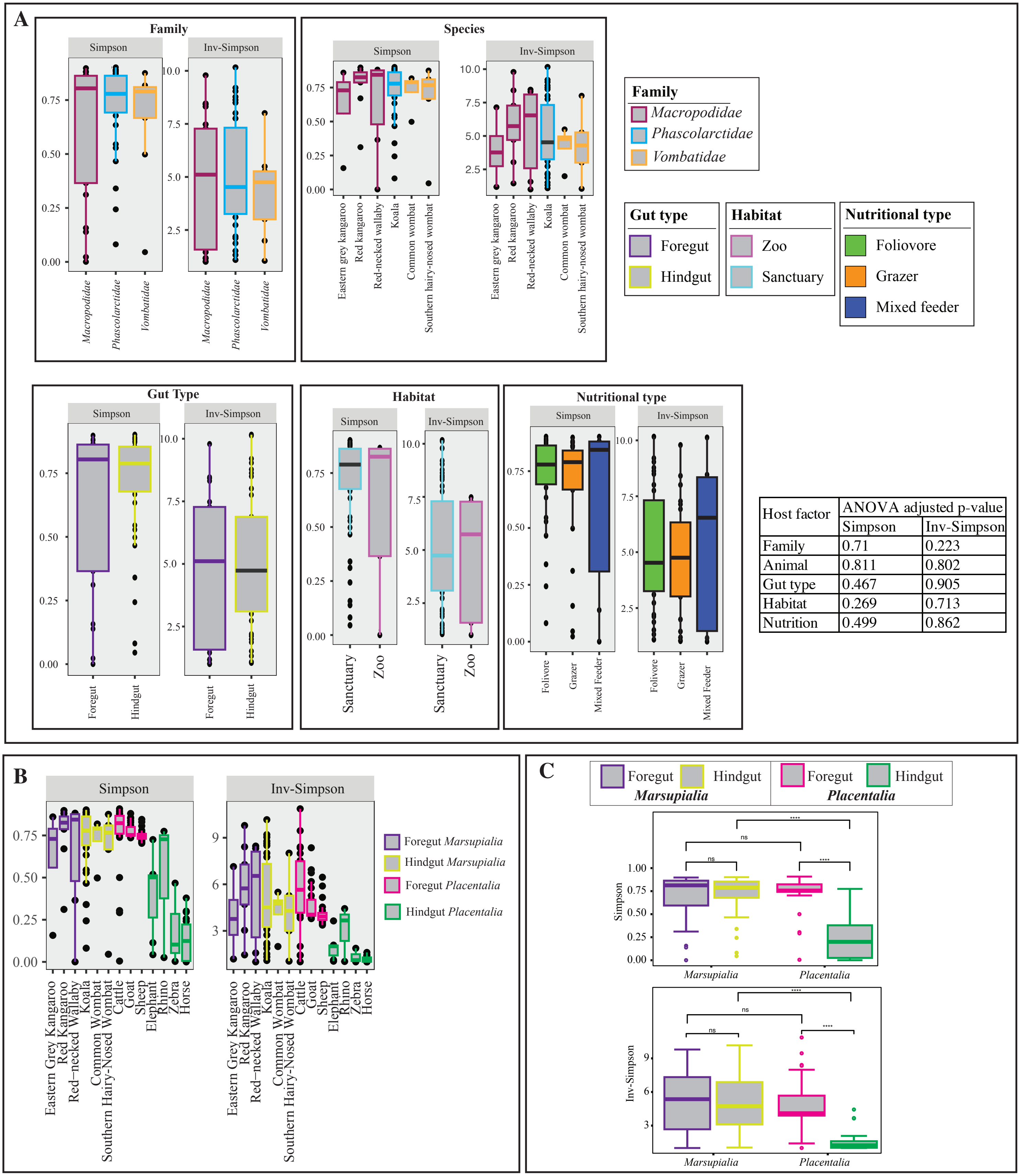


| Host factor | ANOVA adjusted p-value | |
|---|---|---|
| | Simpson | Inv-Simpson |
| Family | 0.71 | 0.223 |
| Animal | 0.811 | 0.802 |
| Gut type | 0.467 | 0.905 |
| Habitat | 0.269 | 0.713 |
| Nutrition | 0.499 | 0.862 |
